# A Novel Natural Killer Cell Expansion Technology for the Development of Cellular Immunotherapies

**DOI:** 10.64898/2026.01.19.700252

**Authors:** Ammelie Svea Boje, Anna Langner, Alexander Jochimsen, Carina Lynn Gehlert, Steffen Krohn, Dorothee Winterberg, Sonja Bendig, Eva Maria Murga Penas, Guranda Chitadze, Monika Brüggemann, Lars Fransecky, Katharina Diemer, Dirk Bauerschlag, Natalie Baum, Daniela Wesch, Hans-Heinrich Oberg, Regina Scherließ, Andreas Günther, Roland Repp, Claudia D Baldus, Thomas Valerius, Friedrich Stölzel, Katja Klausz, Martin Gramatzki, Christian Kellner, Matthias Peipp

## Abstract

Adoptive cell therapy based on Natural Killer (NK) cells holds great promise for the treatment of cancer. For all approaches aiming at utilizing NK cells in immunotherapy, efficient *ex vivo* expansion technologies for the generation on of cytotoxic NK cells are a prerequisite for clinical translation. In this study, a novel multifunctional fusion protein consisting of a CD20-directed Fab-fragment, an agonistic anti-4-1BB single-chain Fragment variable (scFv), the Sushi domain of the interleukin (IL)-15 receptor and human IL-15 was generated. This molecule triggered strong NK cell expansion when bound to co-cultivated autologous B cells, due to trans-presentation of IL-15 and binding to 4-1BB/CD137. Expansion rates of up to 7,500-fold were achieved and the NK cells showed high cytotoxic capacity against a panel of tumor cell lines representing various tumor entities. Importantly, the activated NK cells did not show cytolytic activity against non-malignant B cells indicating that NK cells amplified by our novel approach were still physiologically regulated. The cytotoxic activity of the expanded NK cells was further enhanced by combination with therapeutic antibodies. Our molecule was additionally able to trigger efficient proliferation of NK cells from cord blood as well as multiple myeloma (MM) and acute myeloid leukemia (AML) patients. In conclusion, our novel platform technology provides *ex vivo* expansion of NK cells by using a single multifunctional fusion protein and may be well-suited for the development of NK cell-based immunotherapies.

**Key points:** A novel fusion protein that enables NK cell expansion from different sources including peripheral blood, bone marrow and cord blood

## Introduction

Due to the ability of Natural Killer (NK) cells to recognize and directly kill malignant cells without previous sensitization and their capacity to shape adaptive immune responses by secretion of proinflammatory cytokines, modulating the activity of NK cells *in vivo* or adoptive NK cell transfer represent promising strategies for tumor immunotherapy [1–3]. Different studies have shown that in selected cancer entities NK cells are reduced in number or display an altered phenotype, which plays an important role in tumor development and progression [4–7]. Likewise, for several malignancies it has been shown that the NK cell number, NK cell functionality and the amount of tumor-infiltrating NK cells depict the clinical outcome of different therapies [7, 8]. NK cell activity can be modulated *in vivo* by the application of monoclonal antibodies triggering antibody-dependent cellular cytotoxicity (ADCC) via engagement of Fc gamma receptor (FcγR)IIIa, antibodies blocking critical immune checkpoints such as natural killer group 2 member A (NKG2A) or cytokines to restore immune recognition of tumor cells by NK cells [9–11].

Recently, NK cell-based cellular therapies have gained great attention [3, 12, 13]. NK cells from various sources including peripheral blood from patients or healthy donors, umbilical cord blood, or induced pluripotent stem cells (iPSC) are utilized in clinical applications [14–20]. To enhance their therapeutic potential, NK cells can be activated/polarized e.g., with cytokines such as IL-12, IL-15 or IL-18 to gain a memory-like phenotype or genetically engineered as CAR-NK cells [21–23]. Compared to T cell-based cellular therapies, NK cell therapies may offer advantages such as lower risks of cytokine release syndrome, graft-versus-host disease (GvHD) and suitability for allogeneic off-the-shelf treatments [24, 25]. Ongoing clinical trials are showing promising results, especially in hematologic cancers [26–28]. Due to their comparably short *in vivo* half-life, most protocols in clinical application require high numbers of NK cells and probably repeated dosing to achieve antitumor activity [9, 29]. Therefore, effective technologies for NK cell expansion are necessary. Several approaches have been described, which can roughly be divided into feeder cell-free and feeder cell-based approaches [29, 30]. Feeder cell-free approaches often rely on the use of cytokines that are essential for NK cell development, survival and activation such as IL-2 and IL-15 or cytokine cocktails [31–33], but also synthetic proteins have been described [34, 35]. Feeder cell-based approaches frequently depend on irradiated autologous peripheral blood mononuclear cells (PBMCs) or genetically modified tumor cells such as K562 and rely on costimulatory signals (like CD137) or cell contact-dependent factors [36–39].

Here, we present a novel expansion platform technology based on a multifunctional fusion protein that may simplify the manufacturing of NK cells for adoptive cell therapy. The fusion protein enables NK cell expansion from different sources and the expanded NK cells showed high cytotoxic capacity. Remarkably, non-malignant autologous B cells were not attacked, indicating that the expanded NK cells are physiologically regulated. The cytolytic capacity was further enhanced when the NK cells were combined with therapeutic antibodies. Together, our novel platform technology provides a wide range of possible applications for the development of NK cell-based immunotherapies.

## Material & Methods

### Cell culture

Chinese hamster ovary (CHO)-S cells (FreeStyle™ CHO-S™ Cells) were cultivated in CD CHO medium (+ 1% GlutaMax™,1% HT-Supplement) in a horizontal shaking incubator. After large-scale electroporation-mediated transfection, CHO-S cells were cultivated in CD OptiCHO™ (+ 1% GlutaMax™, 1% PLURONIC™ F-68, 1% HT-Supplement) according to the manufacturer’s recommendations.

Tumor cell lines used for cytotoxicity, cell viability and/or binding assays were obtained from the American Type Culture Collection (ATCC) or the German Collection of Microorganisms and Cell Cultures (DMSZ) (**Supplemental Table 1)**. Cell lines were cultured at 6% atmospheric CO_2_ and 37°C according to the provider’s recommendations and authenticated by STR-analyses and repeatedly monitored for Mycoplasma infection using PCR-based methods.

### Cloning and production of recombinant fusion proteins

The expression vectors coding the fusion proteins RTX-CD137scFv-IL-15; RTX-IL-15; RTX-CD137scFv; HER2-CD137scFv-IL-15 were *de novo* synthesized. A schematic representation of the constructs is provided in **Supplemental Fig. S1A**.

For the production of the fusion proteins, CHO-S cells were transfected by electroporation using the MaxCyte Flow Electroporation^®^ STX Unit and electroporation chamber OC-400 according to the manufacturer’s recommendations. Produced fusion proteins were purified with CaptureSelect™ IgG-CH1 Matrix-based affinity chromatography. Size exclusion chromatography using the ÄKTApure liquid chromatography system (Cytiva) was performed to remove multimers or aggregates.

### Cell viability assay

Functionality of the fused cytokine was evaluated in cell viability assays (Cell Proliferation Kit I, Roche) using the IL-2-dependent cell line CTLL-2. CTLL-2 cells were washed twice in HBSS-buffer and starved for 5 h (withdrawal of IL-2). After starvation, 3×10^4^ cells/well were seeded into 96-well-plates and the respective fusion proteins or recombinant human IL-15 (rIL-15) were added at equimolar concentrations. Cells that did not undergo IL-2 withdrawal served as positive control while cells without the addition of the fusion proteins or cytokines served as negative controls. The cell viability assay was performed according to manufacturer’s recommendations.

### Flow cytometry analyses

All immunofluorescence analyses were performed on a Navios EX-Flow cytometer (Beckman Coulter). For cell surface marker analyses, 2-3×10^5^ cells were incubated with directly labelled primary antibodies (1:20) in the dark for 30 min on ice (**Supplemental Table 2)**. For target cell binding analyses, 3×10^5^ cells were incubated with the respective fusion proteins for 60 min on ice. After the first incubation step, cells were washed thrice with PBS containing 1% BSA and 0.1% sodium azide (PBA) followed by a second incubation step with a respective secondary antibody (anti-human kappa FITC) for 30 min on ice. Cells only stained with the secondary antibody, or corresponding IgG-isotype controls served as negative controls. For CD137 binding analyses, CCRF-CEM cells were stimulated with phorbol 12-myristate 13-acetate (PMA)/ionomycin as described before [40].

### Preparation of PBMCs and isolation of NK and B cells

Citrate anticoagulated blood from healthy donors or patients were obtained and PBMCs were isolated as described [41]. NK cell and B cell isolation was performed with corresponding isolation kits as described by the manufacturer (Miltenyi Biotec). The experiments were approved by the Ethics Committee of the Christian-Albrechts-University of Kiel in accordance with the Declaration of Helsinki.

### NK cell expansion from purified NK cells or patient and cord blood-derived PBMCs

To determine the expansion of NK cells, isolated NK cells were co-cultured with autologous B cells at a ratio of 2:1. Alternatively, freshly isolated patient-derived PBMCs or cord blood samples (Stemcell Technologies) were incubated at a starting condition of 1×10^6^ cells/mL in NK MACS medium (+ 1% NK MACS Supplement, + 5% human AB serum). Co-cultures were cultivated in a 6% CO_2_, 95% atmospheric moisture at 37°C. Fusion proteins were added to the co-cultures at 18.7 nM. After 4 days, media was exchanged and fusion proteins were added. New media and the respective fusion proteins were added every three to four days. The total number of cells was determined and cells were reseeded at 1×10^6^ cells/mL.

### Cytotoxicity assays

The natural cytotoxicity of NK cells or ADCC was evaluated in 4 h ^51^Cr release assays as described before [41, 42]. Isolated NK cells and *ex vivo* expanded NK cells were used as effector cells at effector-to-target cell (E:T) ratios of 10:1 or as stated in the graph. Tumor cell lines from different entities served as target cells. ADCC was evaluated in the presence of monoclonal antibodies. ^51^Cr release was measured by a MicroBeta Trilux 1450 LSC & Luminescence Counter (Perkin Elmer).

### Single cell RNA sequencing

For single-cell RNA sequencing, the BD Rhapsody mRNA Whole Transcriptome Analysis (WTA) Kit was used according to manufacturer’s protocol. Isolated NK cells were sample tagged and directed to library preparation on day 0 and day 20. The library was sequenced on an Illumina NovaSeq 6000 with 2×100bp aiming for 35,000 reads/cell and subsequently aligned with BD Rhapsody Sequence Analysis Pipeline on the Velsera platform. For analysis, the data was normalized and filtered (nFeature_RNA >500 & nFeature_RNA <7000 & percent.mt <35) utilizing the Seurat V5 package in R [43]. Median Number of filtered high quality cells per donor were 6629 for day 0 and 9569 for day 20.

### Data processing and statistical analyses

Graphical and statistical analyses were performed using GraphPad PRISM 9.0. Histograms are represented as mean ± standard error of the mean (SEM) for at least three independent experiments. Statistical statements were done by performing two-way analysis of variance (ANOVA) and suitable post hoc-tests, as indicated. Significance was accepted with p <0.05.

## Results

### Generation of a novel multifunctional fusion protein for the expansion of NK cells

IL-15 and CD137 ligand trans-presentation on genetically modified target cells such as K562 allows the expansion of NK cells *ex vivo* for cellular therapy [44]. To obviate the requirement of genetically modified feeder cells, we designed a novel fusion protein allowing NK cell expansion. To provide IL-15 and 4-1BB trans-signaling, the fusion protein RTX-CD137scFv-IL15 consisting of the Fab fragment of the CD20 antibody rituximab (RTX), an agonistic scFv directed against CD137, the Sushi domain of the IL-15 receptor and IL-15 was generated (**Fig. 1A, Supplemental Fig. S1A**). The protein is designed to bind CD20 on autologous B cells and thereby allows triggering the IL-15 receptor and CD137 on neighboring NK cells in trans (**Fig. 1A**). To investigate the contribution of the individual components of the molecule on NK cell activation and expansion, additional protein variants were generated. These included RTX-IL-15 and RTX-CD137scFv, which lacked individual building blocks of RTX-CD137scFv-IL-15, as well as HER2-CD137scFv-IL-15 in which the RTX Fab was replaced by a HER2-directed Fab (**Supplemental Fig. S1**). The molecular mass and the purity of the fusion proteins were analyzed via SDS-PAGE under non-reducing and reducing conditions (**Supplemental Fig. S1B**) and showed the expected results.

**Figure 1.**
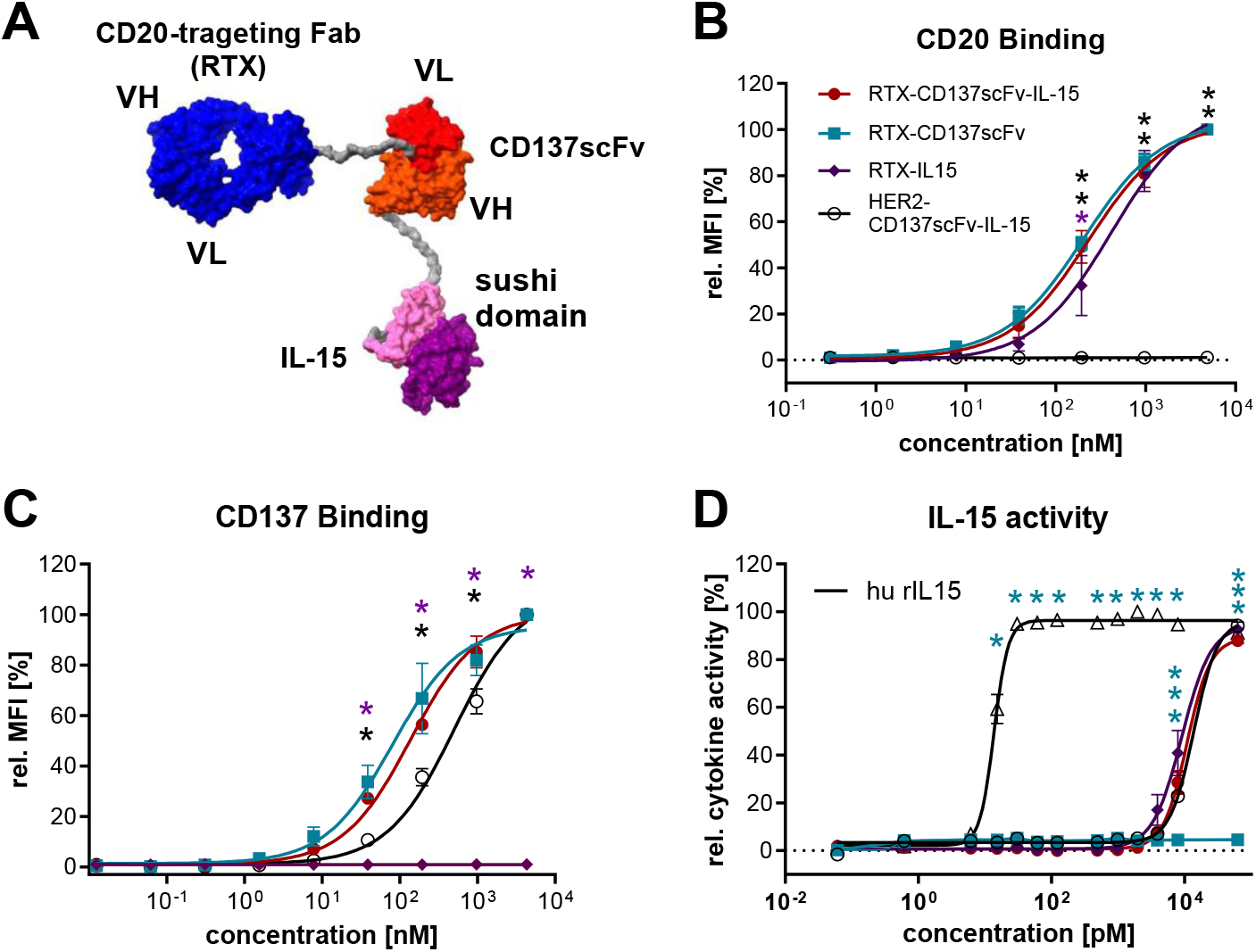
Design and biochemical characterization of the novel NK cell expansion molecule RTX-CD137scFv-IL-15. **A)** Homology model of the RTX-CD137scFv-IL-15 fusion protein; The model of the RTX-CD137scFv-IL-15 fusion protein was generated with MODELLER and Chimera X software using published structures of the rituximab Fab CD20 complex (6VJA), the scFv (6TCS) and the IL-15/IL-15Rα complex (2Z3Q) as templates [67–71]. Fab, fragment antigen binding; scFv, single-chain variable fragment; IL-15, interleukin-15: VH, variable heavy; VL; variable light. **B and C)** Specific antigen binding. Dose-dependent binding of the fusion proteins (RTX-CD137scFv-IL-15 - red filled circle; RTX-CD137scFv - turquoise filled square; RTX-IL-15 - lilac filled diamond; HER2-CD137scFv-IL-15 - black circle) to **B)** CD20^+^ Ramos cells and **C)** 4-1BB/CD137^+^ CCRF-CEM cells. The mean fluorescence intensity (MFI) at saturating concentrations for each cell line were set to 100% and all other experimental values were normalized to this value. Data are presented as mean values of three independent experiments with error bars representing ± SEM, RTX-CD137scFv-IL-15 vs. HER2-CD137scFv-IL-15 and RTX-IL-15; *p≤0.05. **D)** To determine the functionality of IL-15 of the different fusion proteins (RTX-CD137scFv-IL-15 - red filled circle; RTX-CD137scFv - turquoise filled square; RTX-IL-15 - lilac filled diamond; HER2-CD137scFv-IL-15 - black circle), serial dilutions of the fusion proteins were incubated with CTLL-2 cells. Human rIL-15 (black open triangle) served as a control. The mean fluorescence values at saturating concentrations for IL-15 concentration were set to 100% and all other experimental values were normalized to this value. Data are presented as mean values of four independent experiments with error bars representing ± SEM; fusion proteins vs. rIL-15; *p≤0.05, two-way ANOVA.

### RTX Fab and CD137scFv components of the novel RTX-CD137scFv-IL-15 multifunctional fusion proteins show binding to the respective antigens

To determine the binding ability of the fusion proteins, flow cytometric analyses were performed with CD20^+^ Ramos cells (**Fig. 1B**) or PMA/ionomycin-stimulated CD137^+^ CCRF-CEM cells (**Fig. 1C**). The CD20 binding capacity of the CD20-specific fusion proteins (RTX-CD137scFv-IL-15, EC_50_ = 231 nM; RTX-CD137scFv, EC_50_ = 205 nM; RTX-IL-15, EC_50_ = 414 nM) showed no significant differences. No binding on HER2^−^ Ramos cells was observed for the HER2-specific control molecule (**Fig. 1B**). The CD20-specific fusion proteins containing the CD137-specific scFv showed no significant differences in binding to CD137^+^ CCRF-CEM cells (RTX-CD137scFv-IL-15, EC_50_ = 139 nM; RTX-CD137scFv, EC_50_ = 81 nM). Significant differences between the CD20-specific fusion protein containing the CD137-scFv and the HER2-specific molecule (EC_50_ = 511 nM) was observed, indicating that the specific Fab-fragment used to design the fusion protein may impact the overall activity of the molecule. No binding to CD137 was observed for the fusion protein lacking the CD137-specific scFv (**Fig. 1C**).

### The fused IL-15 component of the fusion proteins is functional

To determine the functionality of the IL-15 component of the fusion proteins, a cell viability assay was performed using the IL-15-responsive murine CTLL-2 cell line and rIL-15 as a positive control (**Fig. 1D**). While all fusion proteins containing IL-15 showed stimulatory activity, no significant differences were seen between the fusion proteins. As expected, no stimulatory activity was observed for the fusion protein lacking the IL-15 component. (**Fig. 1D**). Significant differences between rIL-15 and the fusion proteins were observed. While the EC_50_ value of rIL-15 was 0.014 nM, the fusion proteins containing IL-15 showed EC_50_ values between 9 nM (RTX-IL-15) and 14 nM (HER2-CD137scFv-IL-15; **Fig. 1D**). Therefore, the rIL-15 showed an 800-times higher activity compared to the fusion proteins.

### RTX-CD137scFv-IL-15 achieves expansion rates up to 7,500-fold

To determine the capacity of RTX-CD137scFv-IL-15 to trigger NK cell expansion, NK cells were incubated with the fusion protein in the presence of autologous B cells. RTX-CD137scFv-IL-15 potently triggered NK cell expansion with expansion rates between 10 - 7,500-fold after up to 28 days and a mean expansion rate of 2,320-fold (**Fig. 2A**). The purity of the expanded NK cells was investigated after the expansion period (day21-28) and was always greater than 96% (**Fig. 2B**). Importantly, no structural or numerical chromosomal aberrations in the karyotype of the expanded NK cells were detected (**Supplemental Fig. S2**). Furthermore, the requirement of the individual structural components and the necessity of providing IL-15/CD137 signals by trans-presentation to trigger NK cell expansion was analyzed (**Fig. 2C**). In co-cultures of NK cells and B cells only the fusion protein carrying all structural components was able to trigger potent NK cell expansion. In the absence of B cells only minor expansion rates were observed, indicating that trans-presentation was required to provide optimal IL-15/CD137 signaling (**Fig. 2C**).

**Figure 2.**
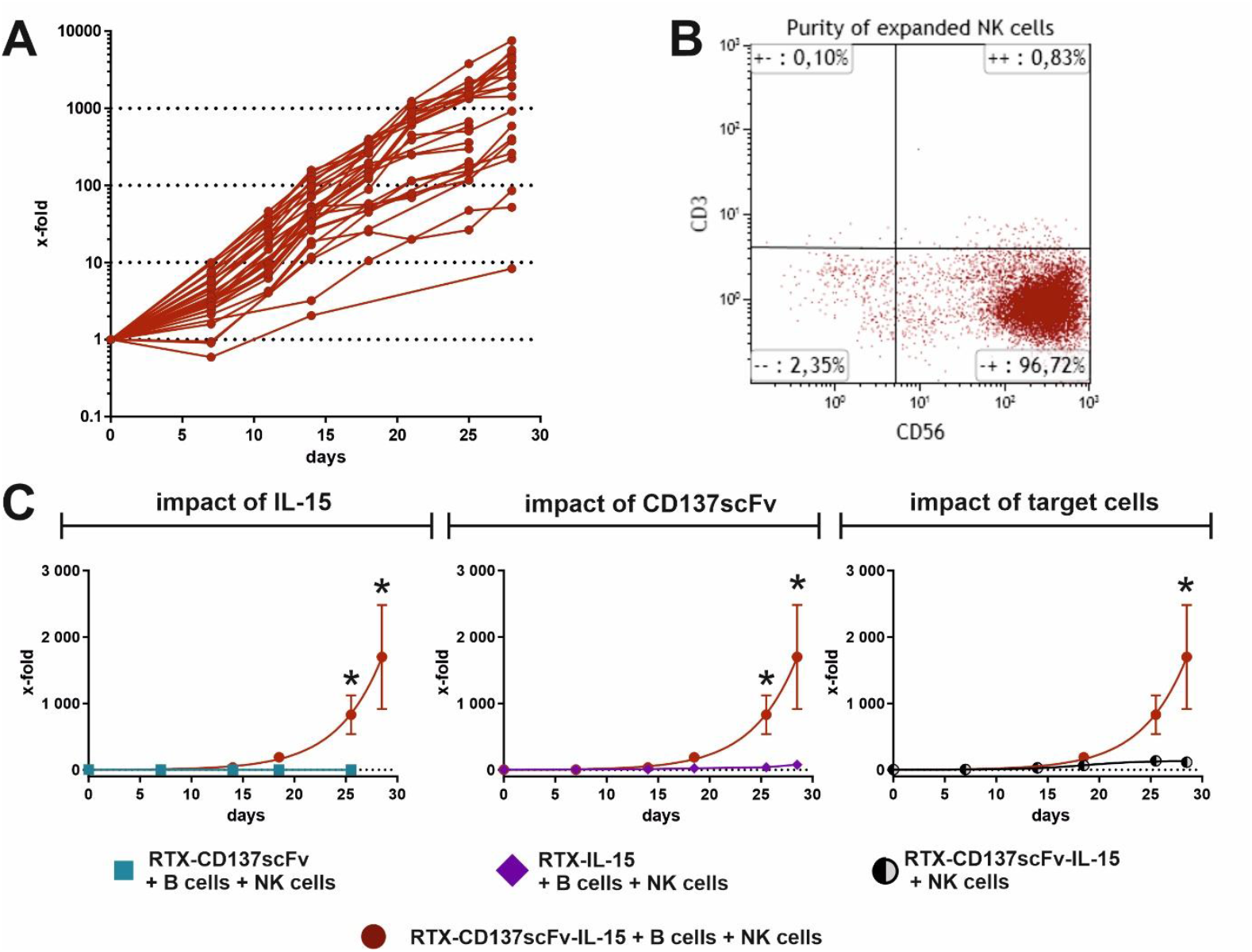
Expansion of NK cells triggered by the novel fusion protein. **A)** Expansion rates of NK cells from 31 different healthy cell donors. The isolated NK cells were co-cultured with autologous B cells and the fusion protein RTX-CD137scFv-IL-15 (18.7 nM) and expanded for up to 28 days. **B)** The purity of the expanded NK cells was checked after the expansion period (day 21-28). Flow cytometric analyses show one representative example. **C)** The need of all structural components was further tested with freshly isolated NK cells from healthy donors co-cultured with autologous B cells and fusion proteins lacking one structural component - either the IL-15 component (RTX-CD137scFv; turquoise square; left graph) or the CD137 component (RTX-IL-15; lilac diamond, middle graph). The impact of target cell presence was tested in expansion experiments where freshly isolated NK cells were incubated with the fusion protein RTX-CD137scFv-IL-15 without the addition of autologous B cells (grey/black circle, right graph). The x-fold expansion is plotted against the time in days. Data are presented as mean values ± SEM of at least three independent donors, *p≤0.05, two-way ANOVA.

### Expansion with RTX-CD137scFv-IL-15 leads to an activated phenotype of the NK cells

To determine the immunophenotype of the expanded NK cells, flow cytometry-based analyses were performed. The expression of different NK cell activation markers and cytotoxicity receptors was determined on day 0 and between days 14-28 of culture (**Fig. 3A**). By comparing the receptor expression at the beginning of the expansion phase with NK cells that underwent expansion, significant changes in the expression of all analyzed activation markers and cytotoxicity receptors was observed (**Fig. 3A**). A significant increase in the expression of activating receptors including NKp30 (3.5-fold), NKp44 (20-fold), NKp46 (1.6-fold), NKG2D (2.5-fold) and DNAM-1 (2-fold) as well as the adhesion molecules CD44 (3-fold) and CD69 (11-fold) was detected. The analyses of the immunophenotype and gene expression profiling experiments confirmed an activated phenotype with upregulation of a panel of cytotoxic mediators. In addition, selected chemokine receptors were upregulated that may impact tissue homing and tissue distribution if applied *in vivo* (**Fig. 3B**). As expected, activation/expansion of NK cells led to upregulation of the inhibitory NKG2A receptor (KLRC1), while other common inhibitory receptors such as LAG3 and TIGIT were only minor or not upregulated (**Fig. 3B**).

**Figure 3.**
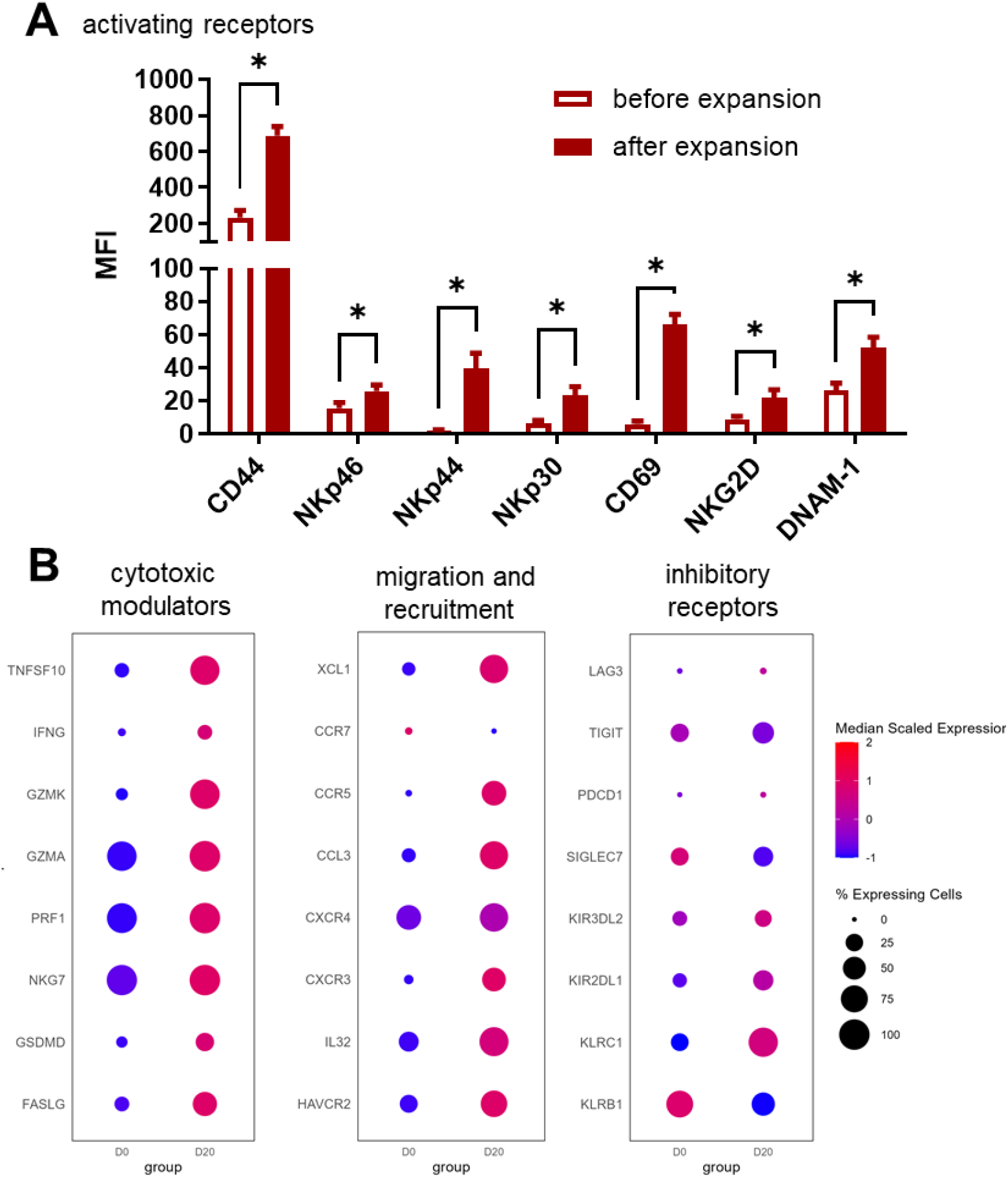
Characterization of *in vitro* expanded NK cells from healthy donors. **A)** The expanded NK cells were characterized by evaluating the expression of different cell surface markers. The activation status of the expanded NK cells as assessed by analyzing a panel of activating NK cell receptors (CD44, NKp46, NKp44, NKp30, CD69, NKG2D and DNAM-1). The expression of these markers was determined by flow cytometer analyses before and after the expansion period. Data are presented as mean values ± SEM of at least five independent NK cell donors, *p≤ 0.05, multiple paired t-test. **B)** Single Cell Gene expression analysis of cytotoxic mediators (TNFSF10 [TRAIL], IFNG, GZMK, GZMA, PRF1, NKG7, GSDMD and FASLG [FAS Ligand]), migration recruitment markers (XCL1, CCR7, CCR5, CCL3, CXCR4, CXCR3, IL32 and HAVCR2 [Tim-3]) and inhibitory and exhaustion genes (LAG3, TIGIT, PDCD1 [PD-1], SIGLEC7, KIR3DL2, KIR2DL1, KLRC1 [NKG2A] and KLRB1) of four independent NK cell donors before expansion (D0) and of the expanded NK cells at day 20 (D20), shown as average expression level across all cells within a class and as median per group.

### Expanded NK cells show high natural cytotoxicity against a panel of tumor cell lines

To determine the cytolytic capacity, NK cells expanded by RTX-CD137scFv-IL-15 were tested in ^51^chromium-release assays with K562 cells as target cells at various effector to target ratios (**Fig. 4A**). The NK cells demonstrated potent lysis, with significant lysis of >30% at a low E:T ratio of 1:1 and >70% at a higher E:T ratio of 20:1. (**Fig. 4A**). Moreover, expanded NK cells were compared to non-stimulated NK cells. Remarkably, expanded NK cells showed a significantly higher natural cytotoxicity against the panel of tumor cell lines compared to non-stimulated NK cells underlining the activated status (**Fig. 4B**). Lastly, natural cytotoxicity of the expanded NK cells was evaluated with a larger panel of tumor cells representing different tumor entities (**Supplemental table 2**). Significant lysis was observed with all tumor cell lines tested ranging from 20% to 75% (**Fig. 4C**). Importantly, no significant lysis was observed with autologous non-malignant B cells, indicating that the highly activated expanded NK cells are still physiologically regulated (**Fig. 4C**).

**Figure 4.**
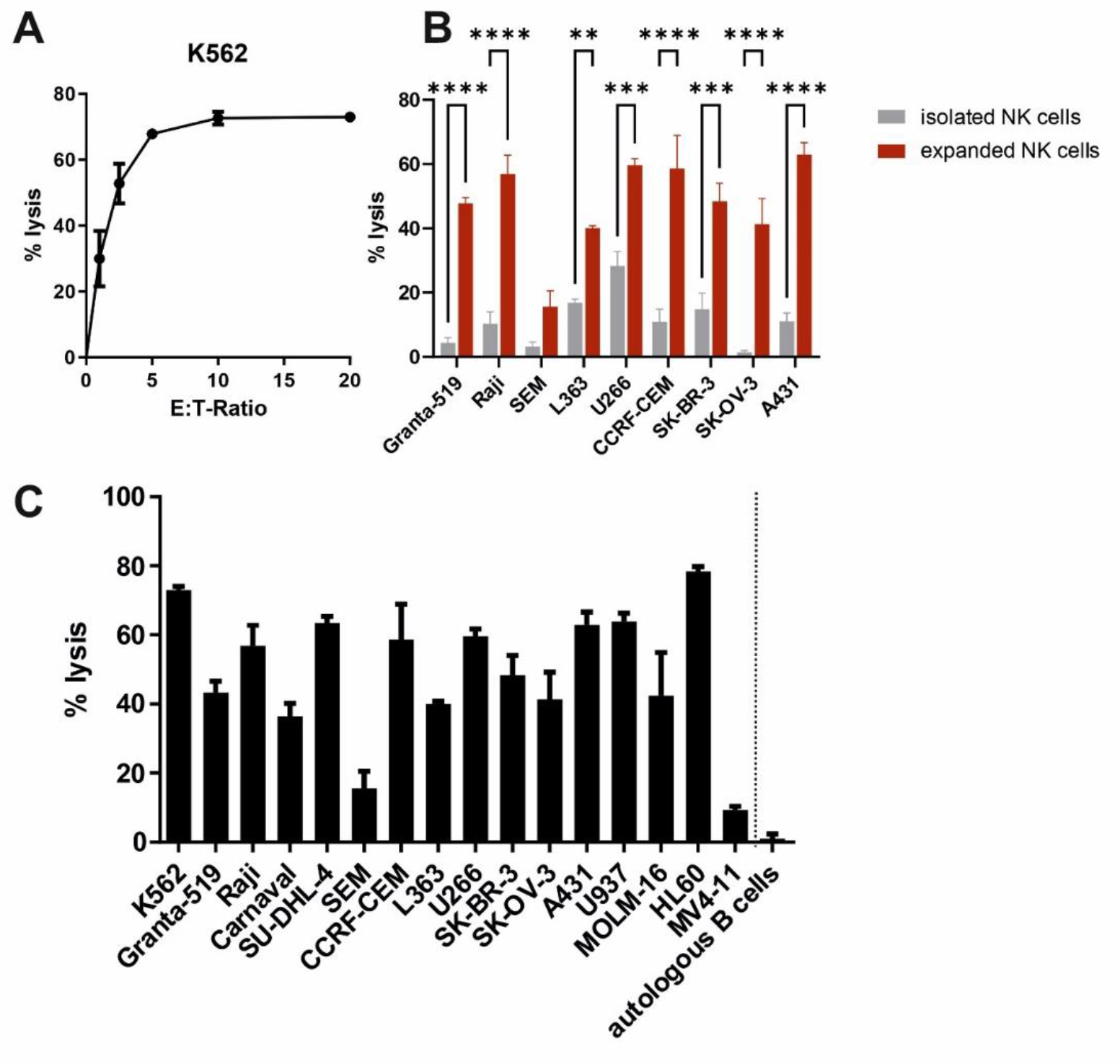
Expanded NK cells show high natural cytotoxicity against a panel of tumor cell lines, but not against autologous non-malignant B cells. **A)** NK cell-mediated tumor cell lysis of K562 cells was tested with expanded NK cells as effector cells at different E:T ratios. **B)** The natural cytotoxicity of expanded NK cells was compared to freshly isolated NK cells in ^51^chromium-release assays with different tumor cell lines as target cells. Both NK cell populations were used at an E:T ratio of 10:1. **C)** The natural cytotoxicity of the NK cells that were expanded with the RTX-CD137scFv-IL-15 expansion molecule was further tested in ^51^chromium release assays with expanded NK cells as effector cells and an extended panel of tumor cells representing different tumor entities or non-malignant B cells. Expanded NK cells were used at an E:T ratio of 10:1. Data presented are mean values ± SEM of three NK cell donors; *p≤ 0.05, **p≤ 0.01, ***p≤ 0.001, ****p≤ 0.0001, two-way ANOVA with Šídák’s-test.

### Cytotoxicity of the expanded NK cells could be enhanced by combination with therapeutic antibodies

Besides natural cytotoxicity, NK cells are capable of triggering ADCC by engagement of FcγRIIIa (CD16a). Flow cytometric analyses revealed that up to 90% of the expanded NK cells showed FcγRIIIa expression (**Fig. 5A**). The capability to trigger ADCC was analyzed with expanded NK cells as effector cells and a panel of hematological and solid tumor cell lines (Granta-519, SEM, L363, SK-OV-3 and A431) and autologous non-malignant B cells as target cells (**Fig. 5B**). Approved antibodies (as mentioned in **Fig. Legend 5C**) were added to compare the natural cytotoxicity to ADCC. Remarkably, the extent of tumor cell lysis was further enhanced by adding a tumor cell-targeting antibody (**Fig. 5B**). Interestingly, by adding rituximab the inhibitory signals and lack of activation signals on non-malignant B cells could be overcome and significant lysis could be achieved (**Fig. 5B**). To evaluate ADCC in more detail, ^51^chromium-release assays were performed with expanded NK cells as effector cells and the CD20^+^ lymphoma cell line Raji as target cells (**Fig. 5C**). The expanded NK cells lysed target cells in the absence of a therapeutic antibody at varying E:T ratios with lysis rates up to 30%. The tumor cell lysis was significantly enhanced up to 60% by adding rituximab. To investigate ADCC in an autologous setting, expanded NK cells were tested in combination with autologous non-malignant B cells as target cells at varying E:T ratios (**Fig. 5D**). No lysis of autologous non-malignant B cells was observed underlining that the expanded NK cells are still physiologically controlled and do not attack non-malignant cells. As shown before, the addition of rituximab lead to significant lysis (**Fig. 5D**).

**Figure 5.**
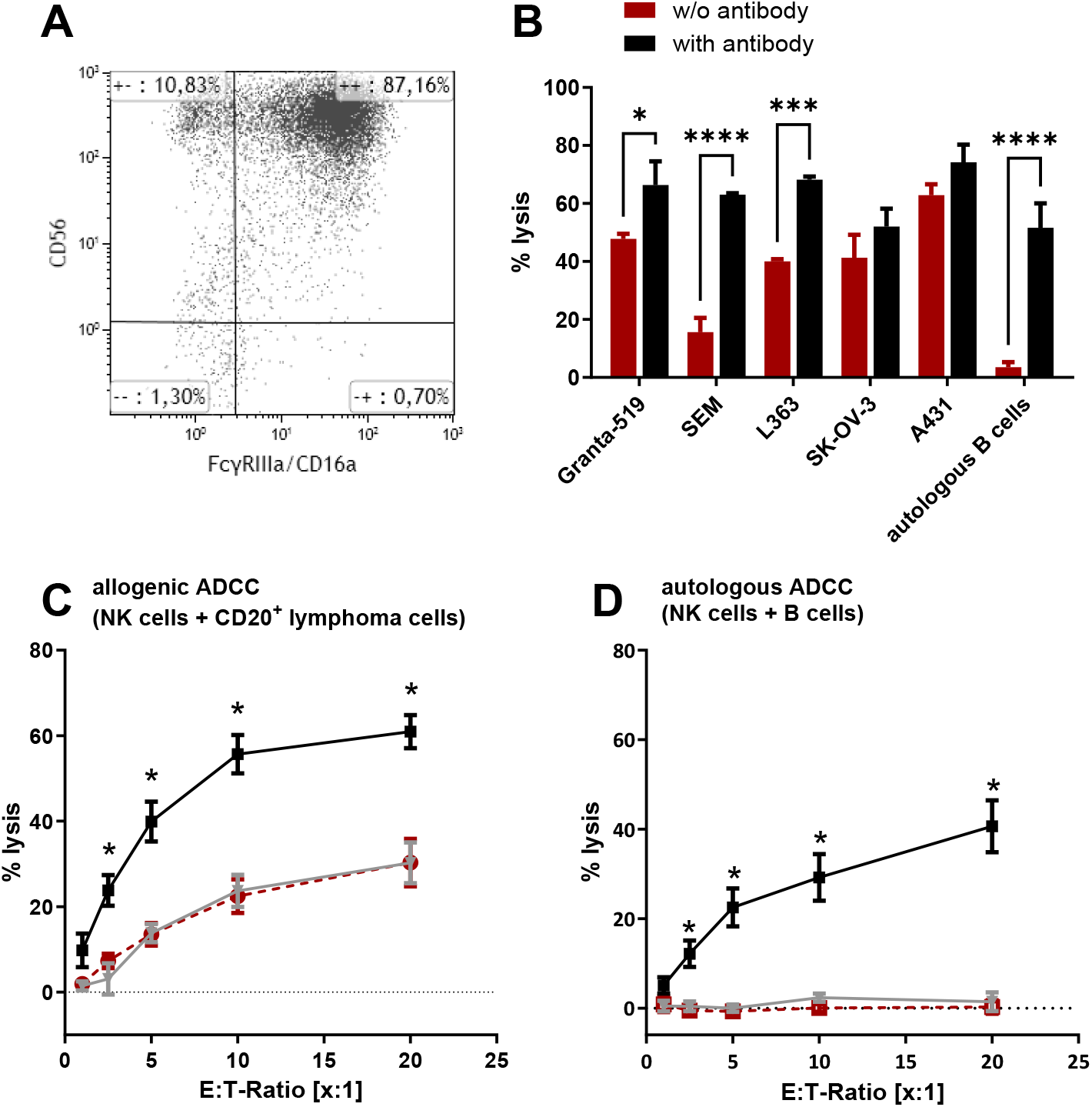
ADCC mediated by expanded NK cells against tumor cells and non-malignant B cells. **A**) Expression of FcγRIIIa/CD16a on expanded NK cells. Flow cytometric analyses show one representative example. **B)** A comparison between the natural cytotoxicity (red bars) and the ADCC (black bars) of the expanded NK cells was tested for different tumor cells lines representing different entities. Tumor cell lysis was determined by performing 4 h ^51^chromium-release assays with expanded NK cells as effector cells and different tumor cell lines (Granta-519, SEM, L363, SK-OV-3 and A431) as target cell lines at an E:T ratio of 10:1. Target cell lysis of autologous non-malignant B cells was also determined. Natural cytotoxicity was measured for effector and target cells without the addition of antibodies. ADCC was quantified for effector and target cells with the addition of the respective antibodies (all with 5µg/mL; Granta-519 - rituximab, SEM - tafasitamab, L363 - daratumumab, SK-OV-3 - trastuzumab, A431 - cetuximab, autologous B cells - rituximab). **C** and **D)** Tumor cell lysis of a CD20^+^ tumor cell line **(C)** and target cell lysis of autologous non-malignant B cells **(D)** through expanded NK cells was tested in 4 h ^51^chromium-release assays. The expanded NK cells were used at different (E:T) cell ratios. Natural cytotoxicity (red dotted line) was compared to ADCC by a respective antibody (rituximab, black line) and a control antibody (trastuzumab, grey line). Data represent mean values ± SEM of three independent NK cell donors; *p≤ 0.05, **p≤ 0.01, ***p≤ 0.001, ****p≤ 0.0001, analyzed by two-way ANOVA.

### RTX-CD137scFv-IL-15 triggers effective expansion of cytotoxically active cord blood-derived NK cells

For clinical application also other sources of NK cells are promising. Therefore, the fusion protein was tested with isolated PBMCs from cord blood. A medium expansion rate of 2,200-fold was achieved ranging between 590-fold and 6,636-fold (**Fig. 6A**). NK cell-mediated tumor cell lysis of K562 cells of up to 70% was detected at an E:T ratio of 20:1 (**Fig. 6B**). Even at lower E:T ratios lysis rates up to 50% were observed indicating that the expanded cord blood-derived NK cells possess a high cytotoxic capacity. This was further evaluated with a larger panel of tumor cell lines representing different entities (**Fig. 6C**). High natural cytotoxicity against all tested tumor cell lines was observed with lysis rates ranging between 70% for K562 and acute myeloid leukemia (AML) cell line HL-60 and 15% for B-cell precursor leukemia SEM (**Fig. 6C**).

**Figure 6.**
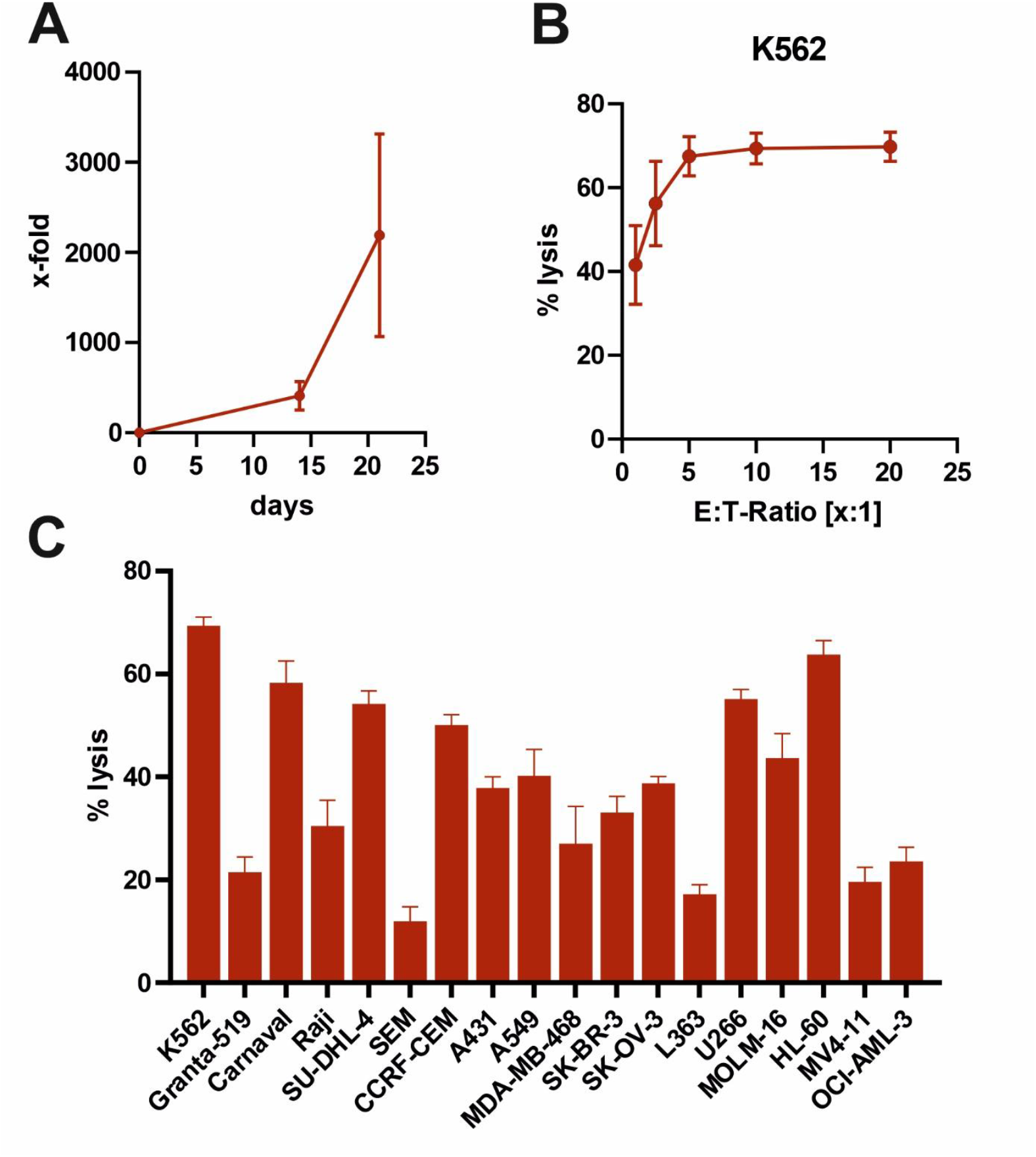
RTX-CD137scv-IL-15 triggers effective expansion of NK cells from cord blood. **A)** As an additional source of NK cells, cord blood-derived PBMCs (n = 5) were cultivated with the novel expansion molecule RTX-CD137scFv-IL-15. The x-fold expansion was plotted against the time in days. **B)** The expanded NK cells (n = 5) showed a high natural cytotoxicity against the tumor cell line K562 in standard ^51^chromium-release assays. The expanded NK cells were used as effector cells against K562 as target cells at different E:T ratios. **C)** The natural cytotoxicity was further tested with tumor cell lines from different entities. ^51^Chromium-release assays were performed with the expanded NK cells as effector cells at E:T ratios of 10:1. Data represent mean values ± SEM of 3-5 cord blood samples.

### RTX-CD137scFv-IL-15 triggers NK cell expansion of cytotoxically active NK cells from multiple myeloma (MM) and AML patients to a high extent

Lastly, we tested whether our novel fusion protein was capable in triggering expansion of NK cells from MM and AML patients (**Supplemental Table 3 and 4**). PBMCs isolated from peripheral blood or bone marrow aspirates of MM and AML patients were used. Interestingly, expansion rates between 10-530-fold were achieved for MM patients (**Fig. 7A, Supplemental Fig. S3**) and 25-14,692-fold for AML patients (**Fig. 7C, Supplemental Fig. S3**). NK cells that were expanded from MM patients were able to lyse MM cell line L363 up to 35%. Moreover, an increased tumor cell killing capacity was observed when the expanded NK cells were combined with the approved CD38 antibody daratumumab (**Fig. 7B**). Additionally, NK cell-dependent tumor cell lysis of autologous MM cells was observed when the expanded NK cells were co-cultured with initially isolated patient tumor cells. The extent of target cell lysis was further enhanced by the addition of daratumumab (**Fig. 7B**). NK cells expanded from AML patients showed a high cytotoxic potential with lysis rates between 20 and 80% when the expanded NK cells were used as effector cells against AML cell lines MOLM-16, HL-60 and MV4-11 (**Fig. 7D**).

**Figure 7.**
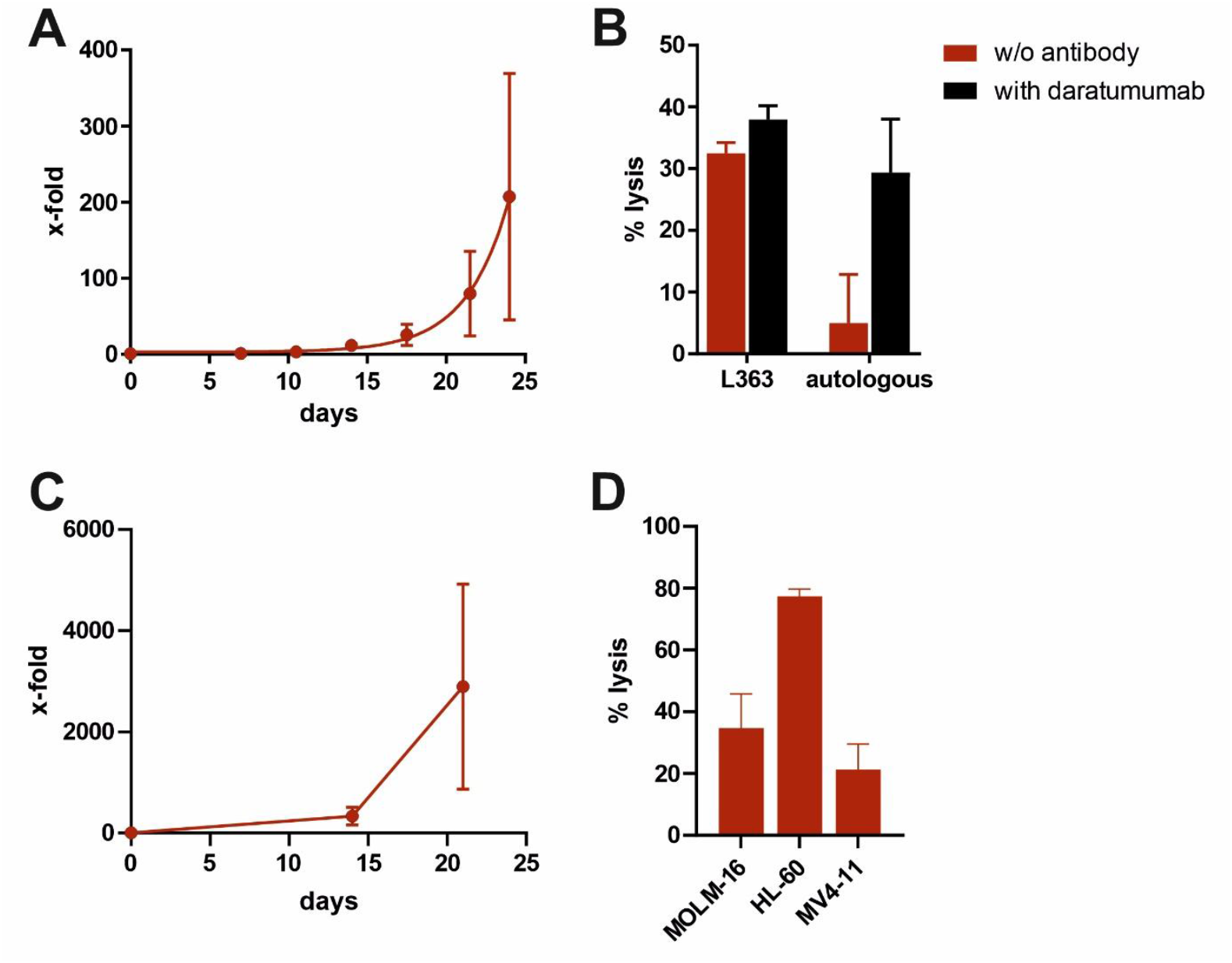
RTX-CD137scFv-IL-15 initiates expansion of NK cells from MM and AML patients. **A** and **C)** Freshly isolated PBMC of MM (n = 6) and AML (n = 7) patients, respectively, were incubated with the expansion molecule RTX-CD137scFv-IL-15 for up to 24 days. The x-fold expansion is plotted against the time in days. **B** and **D)** The cytotoxic capacity of the expanded NK cells was tested in 4 h ^51^chromium-release assays. **B)** NK cells expanded from MM patients were tested against the cell line L363 and autologous patient material at an E:T ratio of 10:1 without or with the addition of daratumumab. **D)** NK cells expanded from AML patients were tested against the cell lines MOLM-16, HL-60 and MV4-11 at an E:T ratio of 10:1. Data represents mean values ± SEM of at least three different patients (1 exemplary patient for the autologous MM sample).

## Discussion

In the current study, we show that by using a novel expansion technology, physiologically regulated NK cells with high cytotoxic activity can be generated. In different tumor entities an exhausted NK cell phenotype, NK cell dysfunction or low NK cell numbers have been correlated with poor prognosis [7, 45]. Confirmative studies in selected hematological malignancies showed that an activated NK cell phenotype and a relatively higher number of NK cells correlate with favorable prognosis [7, 46, 47]. Therefore, different approaches are developed to modulate NK cell function *in vivo* or to make use of NK cells for adoptive cell therapy [48]. Efficient expansion technologies are a prerequisite to develop cellular immunotherapies, hence a variety of approaches have been developed [49]. Here, a novel approach for the *ex vivo* expansion of NK cells was established by using a recombinant multifunctional fusion protein. While our fusion protein, which carries both a CD137-scFv and IL-15, showed high expansion rates, the effect was strongly reduced when one of the stimulatory signals was removed. The same effect was observed, when the expansion was set up without autologous B cells. Both observations underline that trans-presentation of IL-15 and stimulation by CD137 are essential to achieve high expansion rates as described by others [50, 51]. Therefore, opsonizing B cells with our fusion protein in part mimics the principle of genetically modified K562 feeder cells expressing membrane-bound IL-15 and 4-1BB ligand for trans-presentation [37, 52]. However, our approach clearly differentiates from widely used K562 feeder cell-based concepts. In general, no genetically engineered allogeneic tumor cells are required, allowing a more simplified production process also from a regulatory perspective. IL-15 and 4-1BB stimulation is provided in trans in the background of autologous non-malignant B cells expressing significant levels of HLA-I molecules and usually lacking activating ligands involved in induced self-recognition. Therefore, NK cell amplification in contrast to K562-based systems may not purely be driven by “missing-self” and “induced-self” recognition. Inhibitory signaling e.g., by HLA-I molecules and HLA-E expressed by B cells may enable the outgrowth of educated NK cell populations with tolerance to self as suggested by Michen et al. [53] and may further reduce the risk of GvHD [54]. Our initial experiments demonstrating a lack of cytotoxic activity against non-malignant autologous B cells indicate that NK cells expanded by our novel approach were still physiologically regulated and probably recognize the respective self-signals [51, 55]. However, adding rituximab led to a cytotoxic activity against these B cells. This implies that the addition of an antibody could compensate a missing NK cell activation in a setting where the tumor cells lack the activating signals or inhibitory signals dominate. With an average expansion rate of 2,320-fold and a maximum expansion rate up to 7,500-fold, the efficacy of our novel molecule was comparable to other previously described systems [29, 49].

The NK cells expanded with our fusion protein showed an activated phenotype, in line with other reports investigating NK cell expansion by triggering IL-15 and 4-1BB signaling [30]. Accordingly, tumor cell lines originating from different entities were more effectively lysed by *ex vivo* expanded NK cells compared to non-stimulated NK cells. Interestingly, a high proportion of up to 90% of the expanded NK cells expressed FcγRIIIa and potently triggered ADCC when combined with clinically approved monoclonal antibodies. Therefore, the described technology is probably well suited to develop NK cell products that can be combined with therapeutic antibodies or the growing number of NK cell-engaging bispecific antibodies [56–58].

Besides NK cells from peripheral blood and iPSCs, it became evident that also cord blood provides a promising source of NK cells used for allogeneic or off-the-shelf purposes [25, 59, 60]. Therefore, different methods have been evaluated to expand cord blood-derived NK cells [61–63]. Remarkably, also our fusion protein was able to expand cord blood-derived NK cells to a high extent, showing the broad applicability of our approach.

Although allogeneic off-the-shelf NK cells are appealing, in various situations also autologous NK cell have interesting applications [19, 64, 65]. Especially in frail patients, the risk of adverse events may be further reduced. Therefore, we investigated if the fusion protein was able to expand NK cells from AML and MM patients. These patients were at different stages of the disease, ranging from newly diagnosed to patients after different lines of treatment. Remarkably, the novel fusion protein was able to expand NK cells from AML and MM patients to a high extent. Moreover, the expanded NK cells were cytotoxically active and triggered ADCC. This suggests the possibility to use our platform technology to expand autologous NK cells for adoptive cell transfer.

As a perspective, a variety of genetic engineering strategies, such as CRISPR/Cas9-mediated knockdown of selected inhibitory receptors or introducing CAR receptors are available that can be combined with our expansion technology to further enhance the cytotoxic capacity [66]. In conclusion, we provide a novel approach for *ex vivo* expansion of NK cells by using one recombinant multifunctional fusion protein. Our approach may be well suited for the development of a variety of therapies based on adoptive NK cell transfer.

## Acknowledgements

We gratefully acknowledge Anja Muskulus and Britta von Below for excellent technical assistance. MP was supported by the Mildred-Scheel Professorship program by the Deutsche Krebshilfe. and by the Deutsche Forschungsgemeinschaft (German Research Foundation) (project number 444949889) (KFO 5010/1 Clinical Research Unit “CATCH-ALL” to S.B., G.C. and M.B), and through the “Clinician Scientist Program in Evolutionary Medicine” (project number 413490537 to G.C).

## Authorship Contributions

ASB and AL designed and performed experiments and analyzed data. AL, AJ, CG, SK, EMMP, SB, GC, MB, CB, DB, DW, HO, LF, KD, NB, DW, TV, KK, RS, FS, MG and CK performed research and provided essential reagents or patient samples. MP and CK initiated and designed experiments and supervised the study. All authors discussed, contributed in writing and edited the manuscript. All authors have read and agreed to the published version of the manuscript.

