## Supplemental Material for "A Novel Natural Killer Cell Expansion Technology for the Development of Cellular Immunotherapies"

**Supplemental methods**

**Determination of karyotype of expanded NK cells:** Cytogenetic analyses of R-banded chromosomes of 3 male and 2 female healthy donors were performed as previously described [1]. Karyotypes of at least 23 metaphases were analyzed at the 300-band level and described following the guidelines of the International System for Human Cytogenetic Nomenclature (ISCN 2020) [2]. Digital image acquisition, processing, and evaluation were performed using NEON interface for case and image data software version 1.3 (MetaSystems, Altussheim, Germany).

**Supplemental Figures**

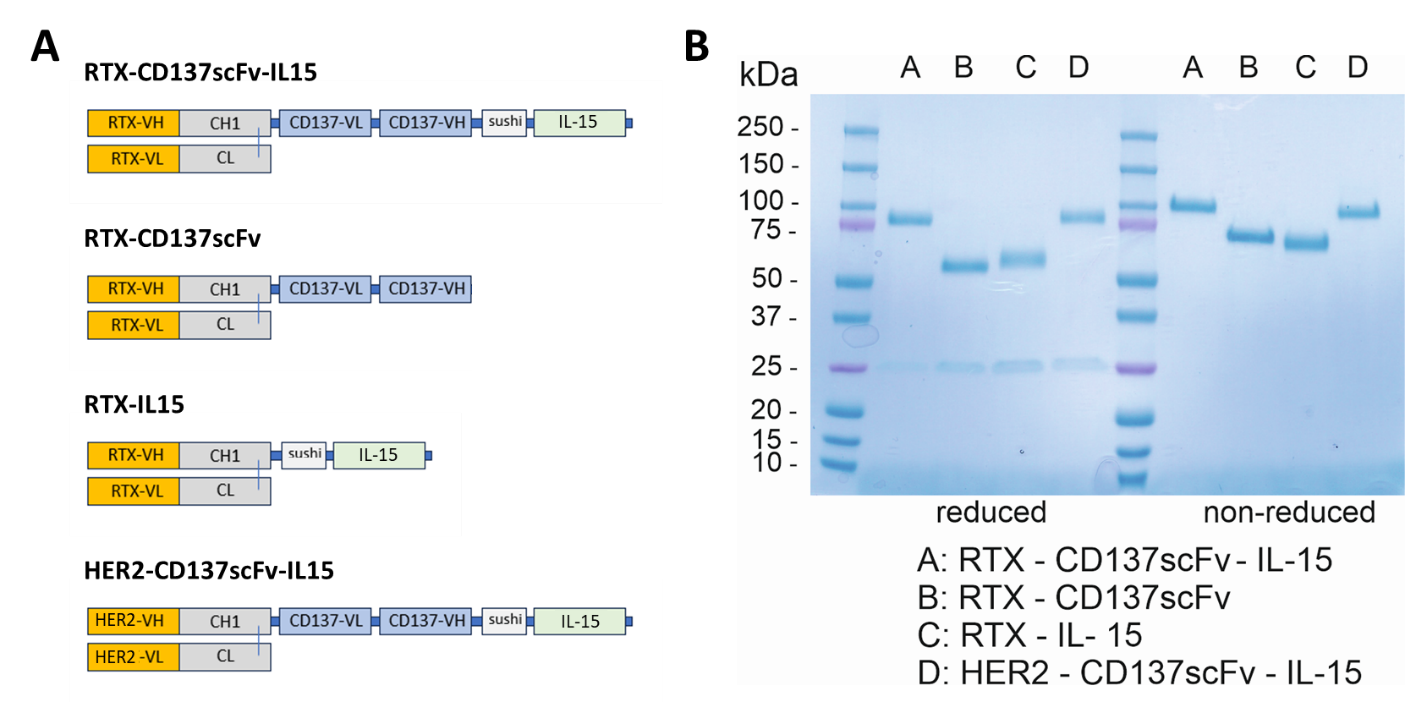

**Supplemental Figure S1.** Schematic overview of the newly generated fusion proteins for the expansion of NK cells and evaluation of purity and size of the fusion molecules by SDS-PAGE and Coomassie staining. **A)** Schematic overview of RTX-CD137scFv-IL-15, RTX-CD137scFv, RTX-IL-15 and HER2-CD137scFv-IL-15 **B)** Two micrograms of purified proteins were analyzed by SDS-PAGE and Coomassie staining using reducing or non-reducing conditions.

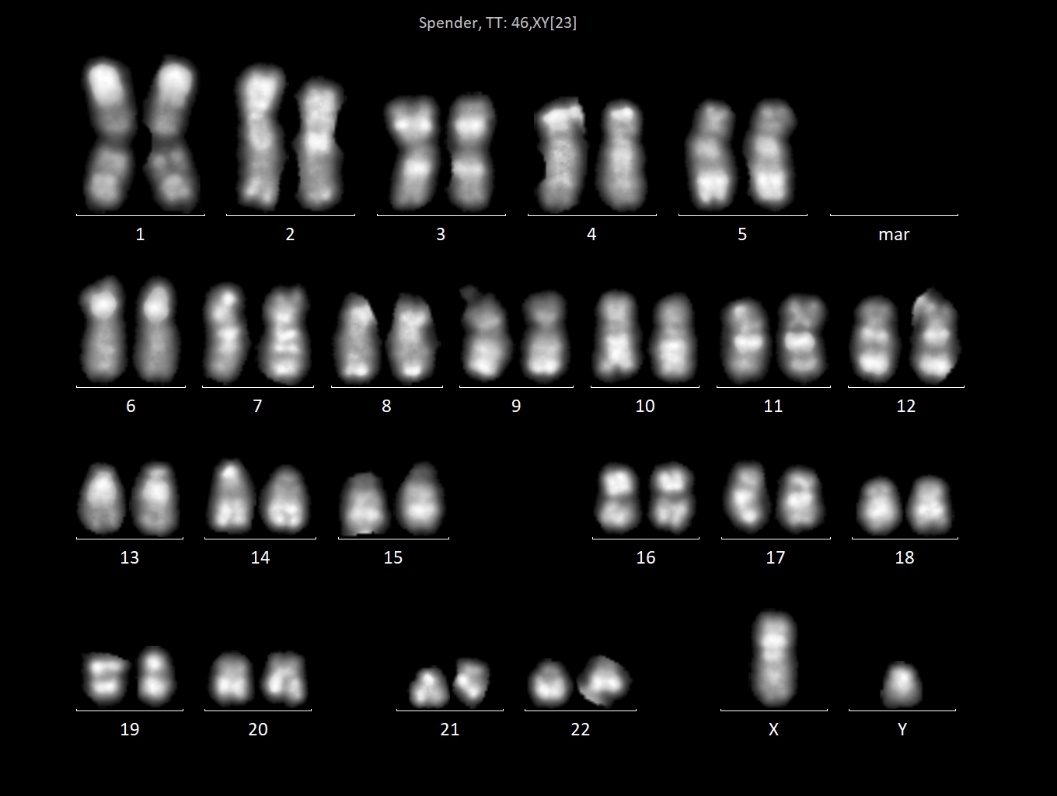

**Supplemental Figure S2. Karyogram of expanded NK cells**. After 20 days of expansion, metaphases of five donors were analyzed. No abnormalities were detected in any of the cases. One representative normal male karyotype is shown.

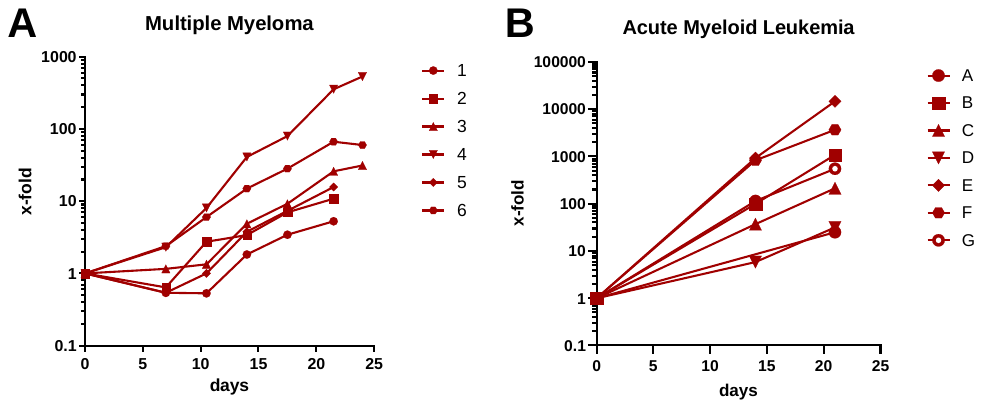

**Supplemental Figure S3. Separated expansion rates of MM and AML patients.** Freshly isolated PBMC of **A)** MM and **B)** AML patients, were incubated with the expansion molecule RTX-CD137scFv-IL-15 for up to 24 days. The x-fold expansion of the individual donors is plotted against the time in days.

**Supplemental tables**

**Supplemental Table 1. Tumor cell lines used** **for cytotoxicity, cell viability and/or binding assays.** Cell lines were cultured in either DMEM (D10^+^), RPMI-1640 (R10^+^) or McCoy´s 5a (McCoy’s20^+^) medium supplemented with 10 or 20% inactivated fetal bovine serum (FBS), 100 U/mL penicillin and 100 µg/mL streptomycin.

| **Tumor cell line** | **Cell line ID** | **Entity** | **media** |
| --- | --- | --- | --- |
| K562 | ACC 10 | chronic myelogenous leukemia | R10^+^ |
| Granta-519 | ACC 342 | B cell Non-Hodgkin lymphoma (B-NHL) | D10^+^ |
| Raji | ACC 319 | Burkitt‘s lymphoma | R10^+^ |
| CARNAVAL | ACC 724 | diffuse large B cell lymphoma (DLBCL) | R10^+^ |
| SU-DHL-4 | ACC 495 | B-NHL | R10^+^ |
| SEM | ACC 546 | B cell precursor acute lymphoblastic leukemia (B-ALL) | R10^+^ |
| CCRF-CEM | ACC 240 | T cell acute lymphoblastic leukemia (T-ALL) | R10^+^ |
| L363 | ACC 49 | plasma cell leukemia (PCL) / multiple myeloma (MM) | R10^+^ |
| U266 | ACC 9 | MM | R10^+^ |
| SK-BR-3 | ACC 736 | breast carcinoma | McCoy´s20^+^ |
| SK-OV-3 | HTB-77 | (ovarian) adenocarcinoma | McCoy´s20^+^ |
| A431 | ACC 91 | epidermoid carcinoma | R10^+^ |
| A549 | ACC 107 | lung carcinoma | D10^+^ |
| MDA-MB-468 | ACC 738 | breast carcinoma | R10^+^ |
| U937 | ACC 5 | monocytic line | R10^+^ |
| MOLM-16 | ACC 555 | acute myeloid leukemia (AML) | R20^+^ |
| HL-60 | ACC 3 | AML | R10^+^ |
| MV4-11 | ACC 102 | AML | R10^+^ |
| OCI-AML-3 | ACC 582 | AML | alpha-MEM10^+^ |
| RAMOS | ACC 603 | Burkitt‘s lymphoma | R20^+^ |
| CTLL-2 | TIB-214 | cytotoxic T lymphocyte (mouse) | R10^+^ + 10% T-STIM with Con A |

**Supplemental Table 2. Antibodies used for flow cytometry analyses**

| **Antibody** | **Antigen** | **Isotype** | **Producer** |
| --- | --- | --- | --- |
| Anti-human-kappa-FITC | Kappa light chain | IgG | Southern Biotech, Birmingham,  AL, USA |
| CD137-PE | CD137 | IgG1 | Beckman Coulter, Brea, CA, USA |
| CD16-FITC | CD16 (FcγRIIIa) | IgG1 | Beckman Coulter, Brea, CA, USA |
| CD19-PE | CD19 | IgG1 | Beckman Coulter, Brea, CA, USA |
| CD20-FITC | CD20 | IgG1 | Beckman Coulter, Brea, CA, USA |
| CD226-PE | CD226 (DNAM-1) | IgG1 | Beckman Coulter, Brea, CA, USA |
| CD314-PE | CD314 (NKG2D) | IgG1 | Beckman Coulter, Brea, CA, USA |
| CD335-PE | CD335 (NKp46) | IgG1 | Beckman Coulter, Brea, CA, USA |
| CD336-PE | CD336 (NKp44) | IgG1 | Beckman Coulter, Brea, CA, USA |
| CD337-PE | CD337 (NKp30) | IgG1 | Beckman Coulter, Brea, CA, USA |
| CD3-CromeO | CD3 | IgG1 | Beckman Coulter, Brea, CA, USA |
| CD44-PE | CD44 | IgG1 | Beckman Coulter, Brea, CA, USA |
| CD56-PC7/PB | CD56 | IgG1 | Beckman Coulter, Brea, CA, USA |
| CD69-PE | CD69 | IgG1 | Beckman Coulter, Brea, CA, USA |
| Isotype-control-FITC | - | IgG1 | Beckman Coulter, Brea, CA, USA |
| Isotype-control-PB | - | IgG1 | Beckman Coulter, Brea, CA, USA |
| Isotype-control-PC7 | - | IgG1 | Beckman Coulter, Brea, CA, USA |
| Isotype-control-PE | - | IgG1 | Beckman Coulter, Brea, CA, USA |
| Isotype-control-CO | - | IgG1 | Beckman Coulter, Brea, CA, USA |

**Supplemental Table 3. Clinical and molecular characteristics of MM patients used in this study**

| **Patient** | **Age** | **Sex** | **Diagnosis** | **M protein** | **previous stem cell transplantation** | **Treatment status** |
| --- | --- | --- | --- | --- | --- | --- |
| 1 | 69 | Male | MM | IgG lambda | none | daratumumab, carfilzomib, lenalidomid, dex |
| 2 | 68 | Male | MM | IgG kappa/light chain kappa | none | untreated |
| 3 | 76 | Male | MM | IgG lambda | auto-SCT | auto-SCT |
| 4 | 60 | Male | MM | IgG kappa | auto-SCT | phase 1 MCL1-inhibitor |
| 5 | 55 | Female | MM | IgG lambda | auto-SCT | lenalidomide |
| 6 | 69 | Female | MM | IgA kappa | allo-SCT | allo-SCT |

**Supplemental Table 4. Clinical and molecular characteristics of AML patients used in this study**

| **Patient** | **Age** | **Sex** | **Diagnosis** | **Relevant mutations** | **Line of therapy** | **Immunosuppression** | **Treatment status** |
| --- | --- | --- | --- | --- | --- | --- | --- |
| A | 74 | Female | AML with TP53 mutation | mTP53 | first line | none | active  Azacitidine |
| B | 64 | Male | AML | del17p | first line | none | Pretherapeutic |
| C | 88 | Male | AML-pCT | mIDH2, mASXL1, mCSFR3, mSTAB2 | first line | MethotrexateGlucocorticoids | active  Venetoclax/  Azacitidine |
| D | 28 | Male | AML with recurrent cytogenetic changes | Molecular aberration | first line | none | active  DA ("7+3"), HDAC |
| E | 37 | Male | AML with KMT2A and SF3B1 mutation | mKMT2AmSF3B1 | second line | none | active |
| F | 71 | Female | AML with monosomy 7 | mRUNX1 VAF, del3q (MECOM) | second line | none | active  Venetoclax/  Azacitidine |
| G | 68 | Male | AML with mutated NPM1 | mNPM1, mIDH1, mDNMT3A, mCEBPA-TAD | first line | none | active  Venetoclax/  Ivosidenib |
